# Population Responses Represent Vocalization Identity, Intensity, and Signal-to-Noise Ratio in Primary Auditory Cortex

**DOI:** 10.1101/2019.12.21.886101

**Authors:** Ruiye Ni, David A. Bender, Dennis L. Barbour

**Affiliations:** Laboratory of Sensory Neuroscience and Neuroengineering Department of Biomedical Engineering Washington University in St. Louis St. Louis, Missouri 63130, U.S.A; Department of Biology Washington University in St. Louis St. Louis, Missouri 63130, U.S.A

**Keywords:** auditory cortex, marmoset monkeys, vocalizations, noise interference, intensity, signal-to-noise ratio, population coding

## Abstract

The ability to process speech signals under challenging listening environments is critical for speech perception. Great efforts have been made to reveal the underlying single unit encoding mechanism. However, big variability is usually discovered in single-unit responses, and the population coding mechanism is yet to be revealed. In this study, we are aimed to study how a population of neurons encodes behaviorally relevant signals subjective to change in intensity and signal-noise-ratio (SNR). We recorded single-unit activity from the primary auditory cortex of awake common marmoset monkeys (Callithrix jacchus) while delivering conspecific vocalizations degraded by two different background noises: broadband white noise (WGN) and vocalization babble (Babble). By pooling all single units together, the pseudo-population analysis showed the population neural responses track intra- and inter-trajectory angle evolutions track vocalization identity and intensity/SNR, respectively. The ability of the trajectory to track the vocalizations attribute was degraded to a different degree by different noises. Discrimination of neural populations evaluated by neural response classifiers revealed that a finer optimal temporal resolution and longer time scale of temporal dynamics were needed for vocalizations in noise than vocalizations at multiple different intensities. The ability of population responses to discriminate between different vocalizations were mostly retained above the detection threshold.

**Significance Statement:** How our brain excels in the challenge of precise acoustic signal encoding against noisy environment is of great interest for scientists. Relatively few studies have strived to tackle this mystery from the perspective of neural population responses. Population analysis reveals the underlying neural encoding mechanism of complex acoustic stimuli based upon a pool of single units via vector coding. We suggest the spatial population response vectors as one important way for neurons to integrate multiple attributes of natural acoustic signals, specifically, marmots’ vocalizations.

## Introduction

Due to the inherent noise in the activity of individual neurons, multiple presentations of an identical sensory stimulus do not yield exactly the same spike trains. By computing the spiking rate averaged across trials to get rid of the response noisiness, researchers have generally expected to estimate the true firing rate driven by a stimulus. Large amounts of single-unit analysis have been conducted in this fashion. In studying neural responses to auditory stimuli, much insight has been gained from analyzing coding properties of individual neurons and at a population level that averaged across individual cells, based upon the simplistic rate-coding hypothesis (Aitkin et al. 1986; Barbour 2011; Bendor and Wang 2005; Imig et al. 1990; Woolley et al. 2006). However, the information represented by the ensemble of individual neurons has typically been overlooked. This seems to be a minor concern for studies investigating relatively simple acoustic stimuli, such as pure tones, but there is study showing that even stationary acoustic stimuli induce dynamic responses on a population level. For more complicated acoustic signals with rich temporal-spectral structures, for instance, marmoset vocalizations (Gehr et al. 2000; Nagarajan et al. 2002), an analytical method for inspecting the response properties among populations is needed.

In contrast to single-unit coding, population coding hypothesizes that the stimulus information is encoded in the brain by a large population of neurons via distributed firing rate patterns (McIlwain 2001). Over the past two decades, multiple population analyses have emerged to reveal the neural encoding and decoding properties at the population level, such as population variability analysis and spatiotemporal coding analysis (Churchland et al. 2010). More importantly, to study the dynamics of population variability in response to vocalizations at multiple intensifies and SNR, this paper further asks whether acoustic features of vocalizations modulate the population variability of the ongoing neural activities.

Neocortical neurons generate time-varying firing patterns with particular temporal structures. In the sensory areas, even presentation of a temporally unstructured stimulus, such as a stationary odor or pure tone, is likely to induce a complex temporal pattern of spiking (Bartho et al. 2009; Stopfer et al. 2003). By unifying the temporally-structured responses of individual neurons, we can visualize the complex spatiotemporal patterns at the population level. The spatiotemporal patterns of the population responses vary with the stimulus when a particular feature of the stimulus is slightly changed, such as intensity (Stopfer et al. 2003). Such visualization analysis has revealed the dynamics of responses to relative simple stimuli, however, little is known about the spatiotemporal patterns of complex vocalization stimuli. Here, by varying intensities and SNR levels, the alteration of spatiotemporal patterns of population neural responses to five marmoset conspecific vocalizations was studied and the hypothesis that the population responses are not just a linear scaling of their amplitude was tested.

Neurometric analysis is a useful tool for linking neural activity with perception to identify the underlying neural substrate that generates the sensory perception and behaviors of interest (Walker et al., 2008). Individual neurons vary widely in their ability to discriminate complex acoustic stimuli (Narayan et al. 2006; Schneider and Woolley 2010; Wang et al. 2007). However, the degree to which a population of neurons can recognize different types of vocalizations at multiple intensities and SNR levels is not well known. Would distributed firing rate patterns across a whole population of recorded cells optimize the performance of the stimulus discrimination task, or is there a subpopulation of neurons that yields the best performance? With respect to the dynamics along the time course, does a population of neurons have a constant discriminability, or are the neural responses at certain time epochs better than at others? These questions were investigated by building population response decoding models that are sensitive to temporal discharge patterns. Using this decoding tool, we further inferred the perception intensity threshold of vocalizations and the detection threshold of vocalizations masked with WGN/Babble noise.

In summary, in this paper, by pooling the activities of individual neurons in response to vocalizations at multiple intensities and SNR levels, we analyzed population responses from three aspects: population response variability with respect to time, the spatiotemporal structures of population responses, and the ability of population responses to identify stimuli under different experimental conditions.

## Materials and Methods

### Surgery and recording

All training, recording, and surgical procedures comply with US National Institute of Health Guide for the Care and Use of Laboratory Animals and are approved by the Washington University in St. Louis Animal Studies Committee.

The experimental materials and procedures were similar to those reported previously (Bender et al. 2016; Ni et al. 2016). Adult common marmoset monkeys (*Callithrix jacchus*) were used as subjects. Subjects were trained to sit upright in a custom, minimally restraining primate chair inside a double walled sound-attenuation booth (IAC 120a-3, Bronx, NY) while their heads were fixed in place by head posts for the same length of duration as that would be for physiology recording. After they became used to this setup, a custom head cap for recording was surgically affixed to the skull of each subject, and temporalis muscle was removed during surgery. The location of the vasculature running within the lateral sulcus was marked on the skull. According to the landmark, microcraniotomies (<1 mm diameter) were drilled through the skull with a custom drill directly over auditory cortex. The location of A1 was identified anatomically based upon lateral sulcus and bregma landmarks and confirmed with physiological mapping (Stephan et al. 2012). Before experiments, the animals were allowed to take sufficient time to recover following the surgery.

A single high-impedance tungsten-epoxy 125 µm electrode (~5 MΩ @ 1 kHz, FHC, Bowdoin, ME) was advanced perpendicularly to the cortical surface within the microcraniotomies. Microelectrode signals were amplified using an AC differential amplifier (AM systems 1800, Sequim, WA) with the differential lead attached to a grounding screw. Single-unit action potentials were sorted online using manual template-based spike-sorting hardware and software (Alpha Omega, Nazareth, Israel). When a template match occurred, the spike-sorting hardware relayed a TTL pulse to DSP system (TDTRX6, Alachua FL) that temporally aligned recorded spike times (2.5 µs accuracy) with stimulus delivery. The recording locations within the head cap were varied daily, covering all the areas of interest.

### Acoustic stimulation

Two types of noises, white Gaussian noise (WGN) and 4-vocalization babble noise (Babble), were mixed with five natural marmoset conspecific-vocalizations from distinct acoustic classes (Trill-Phee, Peep-Trill, Trill-Twitter, Tsik-String and Peep-String) to generate noisy vocalizations at eight different signal-to-noise ratios (SNR; −15 dB SPL to 20 dB SPL with 5dB SPL interval, plus pure noise and clean vocalization). Spectral power was normalized at each SNR level. The five calls were selected to represent most of the acoustic features of a set of twenty calls. In order to distinguish the onset responses induced by the components of noise and vocalization in the synthesized stimuli, two 250 ms intervals of pure noise were concatenated to both ends of each noisy vocalization. Babble noise was created by shuffling superimposed 50 ms-long pieces of four vocalizations, which were different from the five test vocalizations. Both WGN and babble were synthesized with duration equivalent to the longest vocalization, the Trill-Phee. For the other four vocalizations, WGN and babble noise were truncated to the same length as each vocalization.

### Experimental procedure

Single-unit activities in auditory cortex were recorded from two alert adult marmoset monkeys while they were passively listening to the playback of natural and synthesized conspecific vocalizations. Auditory neurons were detected based upon their responses evoked by pure tones and vocalizations. Once an auditory neuron was isolated, its characteristic frequency was estimated by using random spectrum stimuli (Barbour and Wang 2003) and/or pure tones. Next, its rate level functions of five vocalizations were measured with attenuations from 30dB to 90dB with 20dB step in random order with 10 repetitions. The attenuation evoking strong responses to most of the vocalizations was selected to deliver noisy vocalizations. Degraded versions’ of five vocalizations were also randomly displayed with repetitions between 5 and 10.

### Analysis

The dataset is the recorded single-unit responses to five vocalizations (Trillphee, Peeptrill, Trilltwitter, Tsikstring, and Peepstring) at four intensities (from 15 dB SPL to 75 dB SPL, in 20 dB SPL steps). In total, *N* = 326 single units were included in the analysis.

For each unit of both datasets, a peristimulus time histogram (PSTH) for each response trial was calculated by binning spike trains into rate vectors with a 50 ms window in 10 ms steps. The following data analyses are based upon this preprocessing, unless otherwise stated. All data analyses were conducted in MATLAB R2014a (The MathWorks Inc, Natick, MA). Because most of the neurons in our dataset were recorded sequentially one at a time, we created pseudo-populations to substitute for simultaneous recordings. Creating these pseudo-populations potentially ignores the correlation between individual neurons that exists in a simultaneously recorded neuronal population, and may change the estimates of the absolute level of performance. The majority of conclusions drawn in this study, however, would most likely not be altered by the sequential recording, because previous studies show that similar conclusions are obtained by simultaneously recorded neurons and sequentially recorded neurons (Aggelopoulos et al. 2005; Anderson et al. 2007; Baeg et al. 2003; Gochin et al. 1994; Nikolić et al. 2006; Panzeri et al. 2003).

Visualization of population responses in 3D space could help us gain an intuitive understanding about highly complicated neural responses (Bartho et al. 2009; Saha et al. 2013; Stopfer et al. 2003). Such visualization can be realized by principal component analysis (PCA). PCA is a linear dimensionality reduction technique. It identifies a set of linearly uncorrelated variables, called “principal components”, from an original dataset composed of a large number of possibly correlated variables and captures as much of the variability in the dataset as possible (Jolliffe 2002). The principal components are ordered by the amount of variability that each component accounts for, and the first principal component has the largest variance. With regard to neural population responses, each single unit counts as one dimension in the population response space. Given the neural response of *n* single units in a neural population, an n-dimensional response vector *R*^*n*^ can be generated. By applying PCA on the n-dimensional response space, we can obtain m virtual neurons to constitute an m-dimensional response space preserving as much of the variance in the original dataset as possible, where m <= n. If we keep only the first three virtual neurons’ responses (m = 3), we can visualize the population responses as a trajectory in 3D space by connecting the responses at a series of time points. In the following analysis, PCA was implemented for each vocalization type. The first three principal components accounted for about 30% of the original dataset’s variance for each vocalization at multiple intensities, and for about 22% of the variance for each vocalization at multiple SNR levels, under either WGN or Babble noise. For visualization, trajectories were smoothed with a 10-point running window for pure vocalizations, and a 20-point running widow for noisy vocalizations.

Based upon the trajectory visualization analysis, we can further quantify the structure of trial-averaged population responses. To quantify the rotation of the population response vectors in response to a particular stimulus *s*_0_ in 3D space, the angle between a response vector 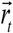 at time *t* and a reference response vector 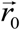 at time *t*_0_ can be computed as in (2) (Bartho et al. 2009). By defining the first point of the spontaneous population response corresponding to stimulus *s*_0_ as the reference vector, the angle evolution of population response vectors can be obtained by concatenating the angles calculated at all the available time points, *t* = *t*_0_, *t*_1_, … *t*_*n*_, during pre-stimulus, stimulus, and post-stimulus. We calculated the angle evolution for the five vocalizations at four intensities and ten SNR levels under WGN and Babble noise conditions as follows:

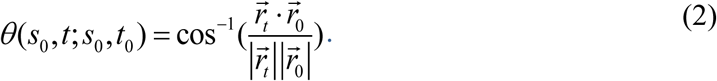

Similarly, we also calculated the angle evolution of population response vectors 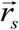 corresponding to stimulus *s* relative to response vector 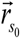, which in turn corresponds to stimulus *s*_0_ across time, *t* = *t*_0_,*t*_1_,…,*t*_*n*_, as displayed in (3).

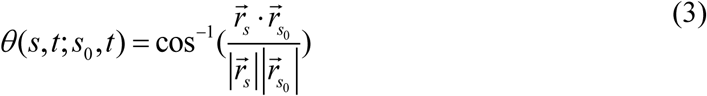

The angles of population responses to vocalizations at three softer (15 dB SPL, 35 dB SPL, and 55 dB SPL) levels relative to the population response to vocalization at 75 dB SPL were calculated. The noise effects on population response angle evolution were also investigated by computing the angles of responses to noisy vocalizations and pure noise relative to pure vocalizations.

To investigate the discriminability of population responses trial-by-trial (Bartho et al. 2009; Meyers et al. 2008), template-based stimulus identity predictive models were built based upon neural population responses. For predictive model decoding of vocalizations at multiple intensities, there are three types of models: single-bin based, sliding-bin based, and varying-cell-number based. To build a single-bin based model, for each stimulus condition (five vocalizations at four intensities), five trials of single-unit responses at a particular time bin were randomly sampled out of, at most, 10 trials for each neuron (N = 326, each has 5~10 trials). The five trials were concatenated to form a 100 × 326 population response matrix. The stimulus identity corresponding to each response trial is called a label. There are a total of five labels, each representing a vocalization type (c = 5). Four trials of the neural population’s response to each of five vocalizations delivered at the highest intensity, 75 dB SPL, were further randomly selected trials as the training templates. The remaining 80 trials of neural population responses were used as the testing dataset, and the label corresponding to each trial was decoded by calculating the cosine distance between this trial and the 20 template trials. The vocalization type or stimulus label corresponding to the template trial with the shortest distance from the test trials was assigned as the predicted label. This whole process was repeated 50 times. The performance of the predictive model was evaluated by its overall accuracy and confusion matrix. The overall accuracy computed the percentage of stimulus labels correctly predicted, and the confusion matrix revealed more information about the chance of a particular label being predicted as one of the five labels.

While a single-bin based model was built to investigate the neural discriminative performance at each time point, a sliding-bin based model was used to study the effect of time accumulation on neural discriminability. The process of building a sliding-bin based model was very similar to that of a single-bin based model, except for that the response bin from each single unit were varied from 1 to 41, where 41 is the number of bins that the shortest vocalization Tsikstring has. Performance as a function of temporal resolutions (time bin width) was investigated with predictive model using 41 bins, and the temporal resolution was varied from 5 ms, 10 ms, 20 ms, …, to 100 ms. Last, a varying-cell-number based model was created by changing the number of neurons in the population, and only models with 41 bins were studied.

The population neural discriminability of vocalizations at multiple SNR levels was investigated, much like that of vocalizations at multiple intensities. Predictive models under WGN and Babble conditions were built separately. Here, the training templates had six labels, including five pure vocalizations and one type of noise (c = 6). In addition, to further study population decoding with a subpopulation of neurons, a predictive model for each vocalization type was built by using the number of time bins available for that particular vocalization. For each vocalization, a different subpopulation of neurons was included because of the contextual dependent effect (Ni et al. 2016). There were only two labels, vocalization and pure noise (c = 2). Population neural responses from ten SNR levels were decoded as either vocalization-present or vocalization-absent. A linear support vector machine (SVM) classifier was used instead of the template-based predictive model to accomplish the binary classification task. The SVM classified the neuronal responses by training a separating hyperplane based upon the labeled training trials and had very good performance for binary classification (Van Gestel et al. 2002), while the template-based method did not have a particular training session. The generalizability of the classifier over lower SNR levels was studied by using different training data, for instance, neural responses to vocalizations at 20 dB SNR.

Normality was verified by the Lilliefors test. Unless otherwise indicated, hypothesis testing was conducted using a two-sided Wilcoxon signed-rank test. The significance criterion was set to 0.05.

## Results

In this study we investigated the alternation of neural population activities evoked by two types of noise, WGN and Babble, in response to marmoset conspecific vocalizations. We recorded the responses of 326 marmoset A1 single-units to five vocalizations degraded by WGN and marmoset Babble, and units were selected for further analysis according to the response criterion for each vocalization. Power spectra of vocalizations and noises are shown in Figure 1A. Spectrograms of vocalization Trillphee at all the tested SNR levels under two noise conditions we studied are also displayed in Figure 1B. For each vocalization, we only included responsive neurons to that particular vocalization in the analysis. To measure the degree of degradation that noise contributed to the neural representation of vocalizations, we calculated the amount of vocalization encoded by each unit as a function of SNR.

**Figure 1:**
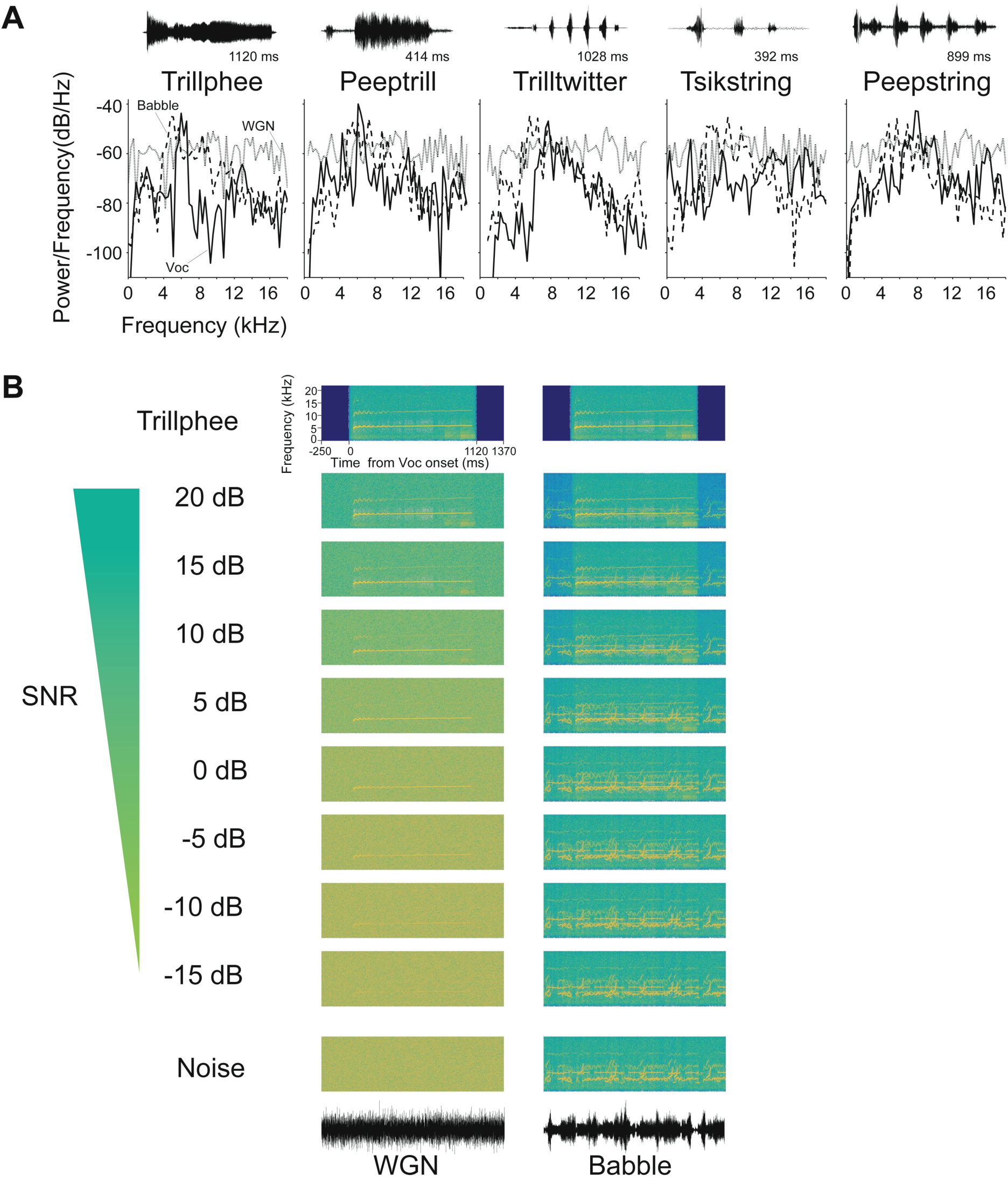
Acoustic stimuli used to investigate robust sound encoding in auditory cortex. (A) Power spectrum of five vocalizations (solid lines), WGN (gray lines) and Babble noise (dashed lines). Background noises were truncated to have the same duration as each vocalization. The temporal waveform of each vocalization is displayed above each power spectrum. (B) Example spectrogram of vocalization Trillphee in noise at 10 different SNR levels, including pure noise and pure vocalization. The first column is Trillphee with WGN as background noise, and the second column is with Babble as background noise. The temporal waveform of WGN and Babble are shown below each column.

### Population coding trajectory for clean vocalizations in 3D response space

An intuitive understanding of population neural responses can be obtained by visualizing their spatiotemporal structure. A powerful tool to achieve this is to project the high-dimensional response vectors onto a lower dimensional space, in which enough variance in the high-dimensional dataset is captured by three principal components (i.e., virtual neurons).

For each vocalization at multiple intensities, population responses were reduced to the same 3D space, and the resulting response trajectories are displayed in Figure 2. Trajectories were formed by connecting the response points from three time stages: pre-stimulus, during-stimulus, and post-stimulus. Skeletons, which link the first point on the trajectory with the remaining points, were plotted to visualize the response hyperplane. Hyperplanes belonging to different vocalizations all have very distinct shapes. Some are relatively smooth and simple, such as Trillphee, while some are more tangled and twisted, such as Peepstring.

**Figure 2:**
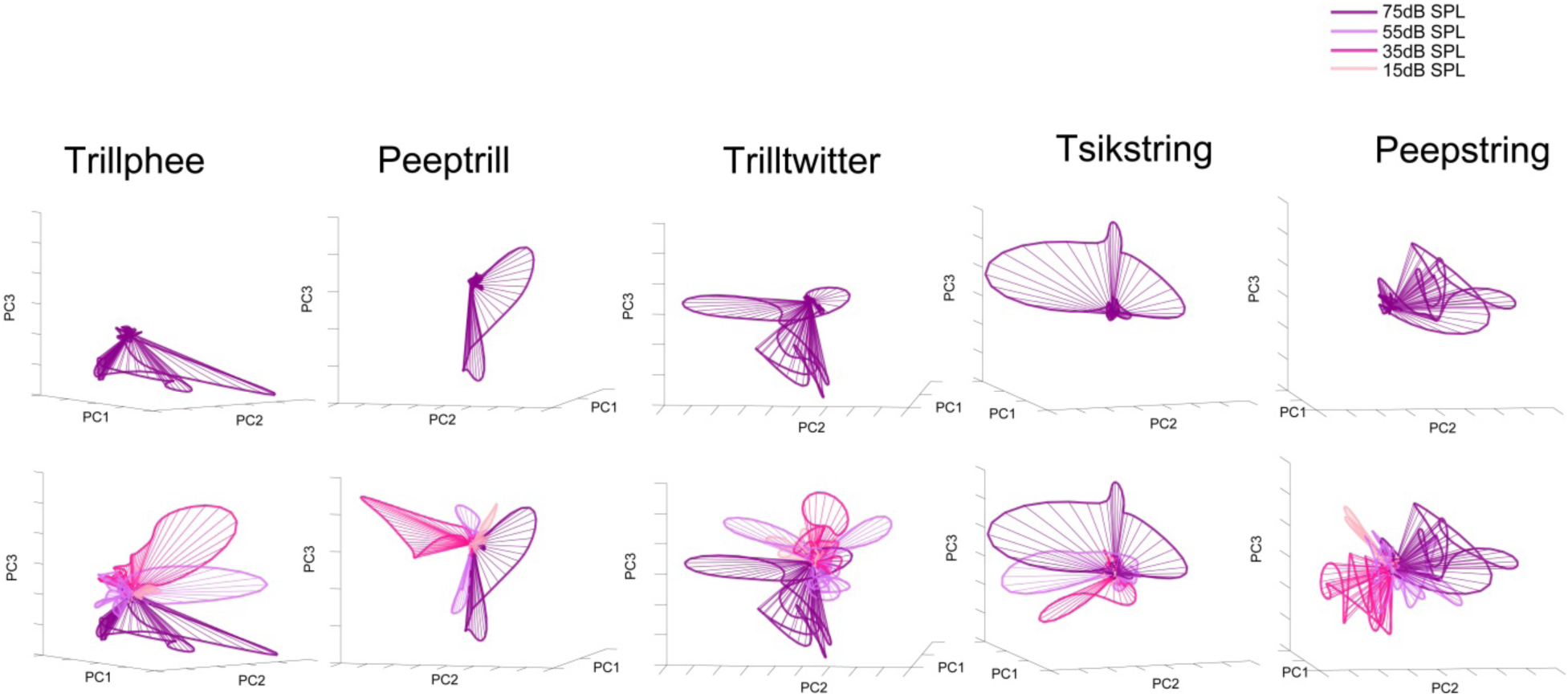
Trajectories of population responses to vocalizations at multiple intensities in 3D space

### Intra-trajectory angle evolution encodes stimulus identity

How does the population hyperplane change in response to a decrease in intensity? Here we consider the hyperplane at 75 dB SPL as the reference hyperplane. If neuronal populations linearly scaled their responses’ amplitudes, we would expect to see the response hyperplane shrink without changing its position in 3D space. Alternatively, the hyperplane could change in a way that only rotates its position relative to the reference hyperplane. As a matter of fact, the hyperplane seems to both resize and rotate. It is worth noting that the more the intensity decreases, the further the hyperplane deviates from the reference, in a consistent direction.

To quantify the response hyperplane and the changed induced by intensity, we calculated the angle between response vectors in two ways. First, we quantitatively described the spatiotemporal structure of a hyperplane by computing the angle between the first response vector on the hyperplane and the remaining response vectors over time in Figure 3. Clearly, across vocalizations and intensities (except for 15 dB SPL), the angle fluctuated between 0 and 60 degrees at the pre-stimulus stage. To process the upcoming stimulus, an acute increase in the angle immediately followed the stimulus onset and further evolved during the stimulus presentation. As the end of the stimulus presentations approached, the angles acutely declined back to the pre-stimulus level. Therefore, the angles of response vectors during the stimulus presentations occupied a distinctively different range from the pre/post presentation. Comparing the angles over time across different intensities, we noticed that at measured intensities above 15 dB SPL, the angles over time were very similar without the scaling shown in the spiking rate and response variability. This similarity indicates that population responses may represent a vocalization identity in an intensity-invariant manner by encoding the information in the angle evolution of a trajectory.

**Figure 3:**
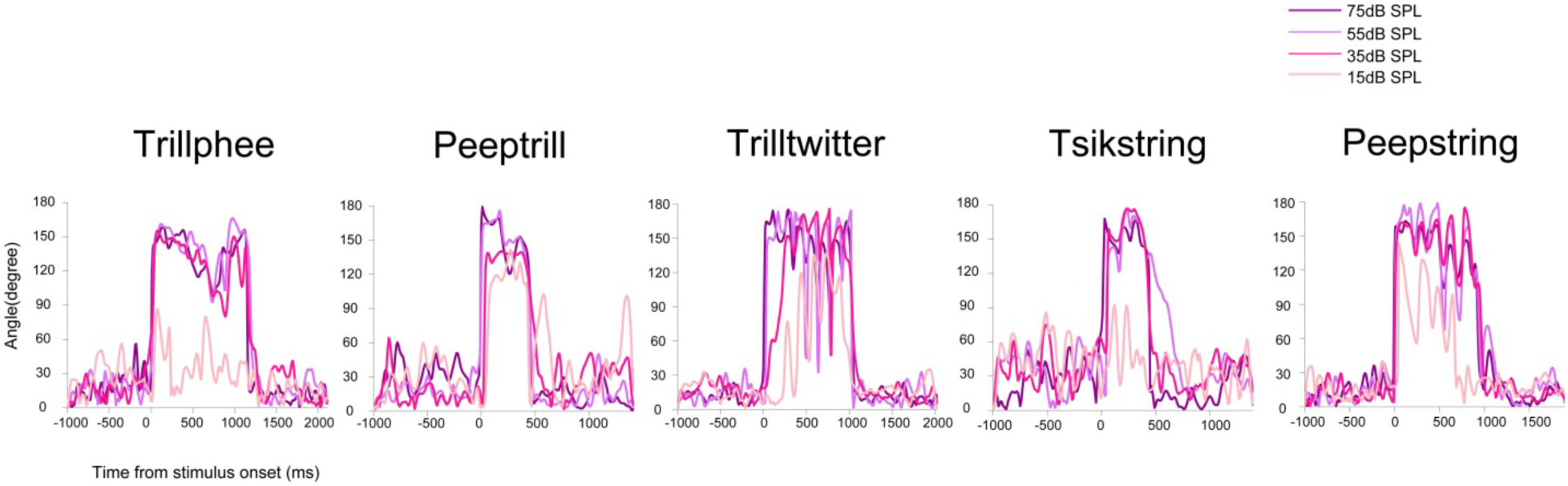
Evolution of rotation angles relative to the first time point (in silence) of the population response at multiple intensities in 3D space

### Inter-trajectory angle evolution encodes stimulus intensity

Next, we quantified the influence of intensity on the deviation of response trajectories by computing the angles between response trajectories of decreasing intensities relative to the reference trajectory at 75 dB SPL over time, as shown in Figure 4. The less intense the vocalization, the further away the corresponding population trajectory was from the reference trajectory in terms of angle rotations, which is consistent with a qualitative visual inspection in Figure 2. The rotation angles, however, are not equal over time. The pre-stimulus and post-stimulus periods have rotation angles that fluctuated in the same range as that in Figure 3. For the stimulus-driven angle evolution, finer structures potentially related to the acoustic features of vocalizations can be observed. With regard to the angles between two neighboring intensities, for instance, 55 dB SPL vs 35 dB SPL, their difference at each time point is generally smaller than the angle difference between different points belonging to the same intensity. Rotation of population responses may serve as an indicator to encode the information of intensity.

**Figure 4:**
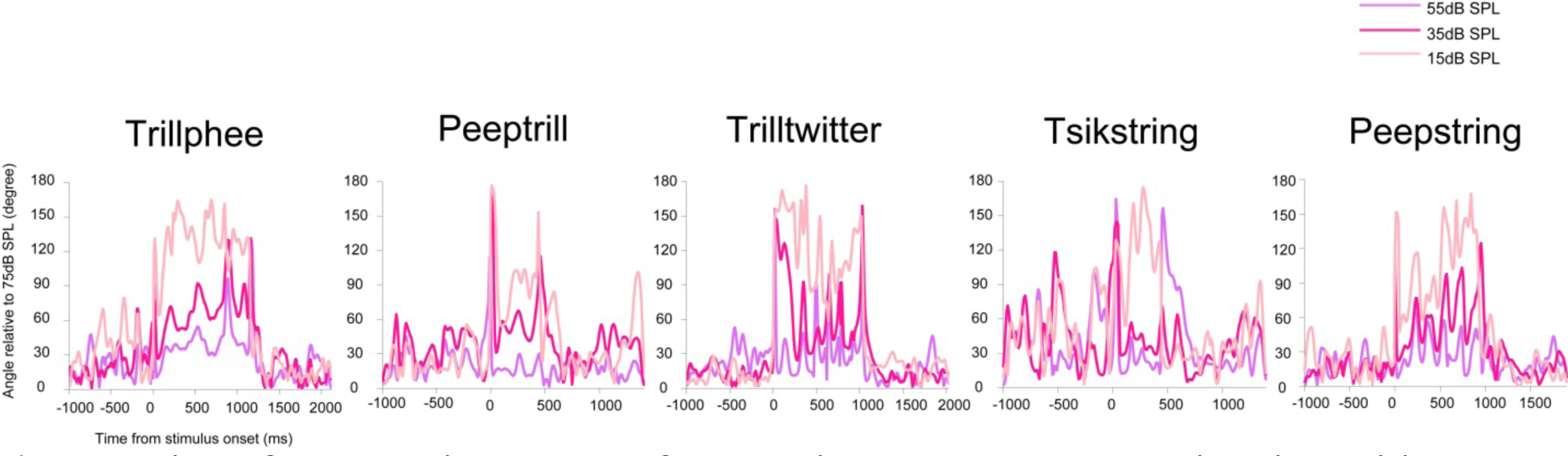
Evolution of the rotation angles of population responses at multiple intensities relative to the population response at 75dB SPL in 3D space

To summarize, population responses to the same vocalization largely retain their intrinsic structures within trajectories in 3D space at multiple intensities. By contrast, the relationship between hyperplanes at different intensities is more complicated than just an equal angle shift.

### Visualization of population coding trajectory for noisy vocalizations in 3D response space

To characterize the spatiotemporal structures of population responses to vocalizations with increasing amounts of noise, the associated population response trajectories based upon three principal components are displayed in Figure 5. Responses were projected to different 3D spaces under two noise conditions, but a salient differences can detected between how increasing the amount of different types of noise affects the population response structures. Under the WGN condition, two groups can be identified. Trajectories to vocalization and 20 dB SNR are clustered together, while trajectories to −10 dB SNR and pure WGN noise share a similar subspace. In contrast, trajectories to vocalizations masked with Babble noise don not form individual clusters, with a large portion overlapped across SNR levels.

**Figure 5:**
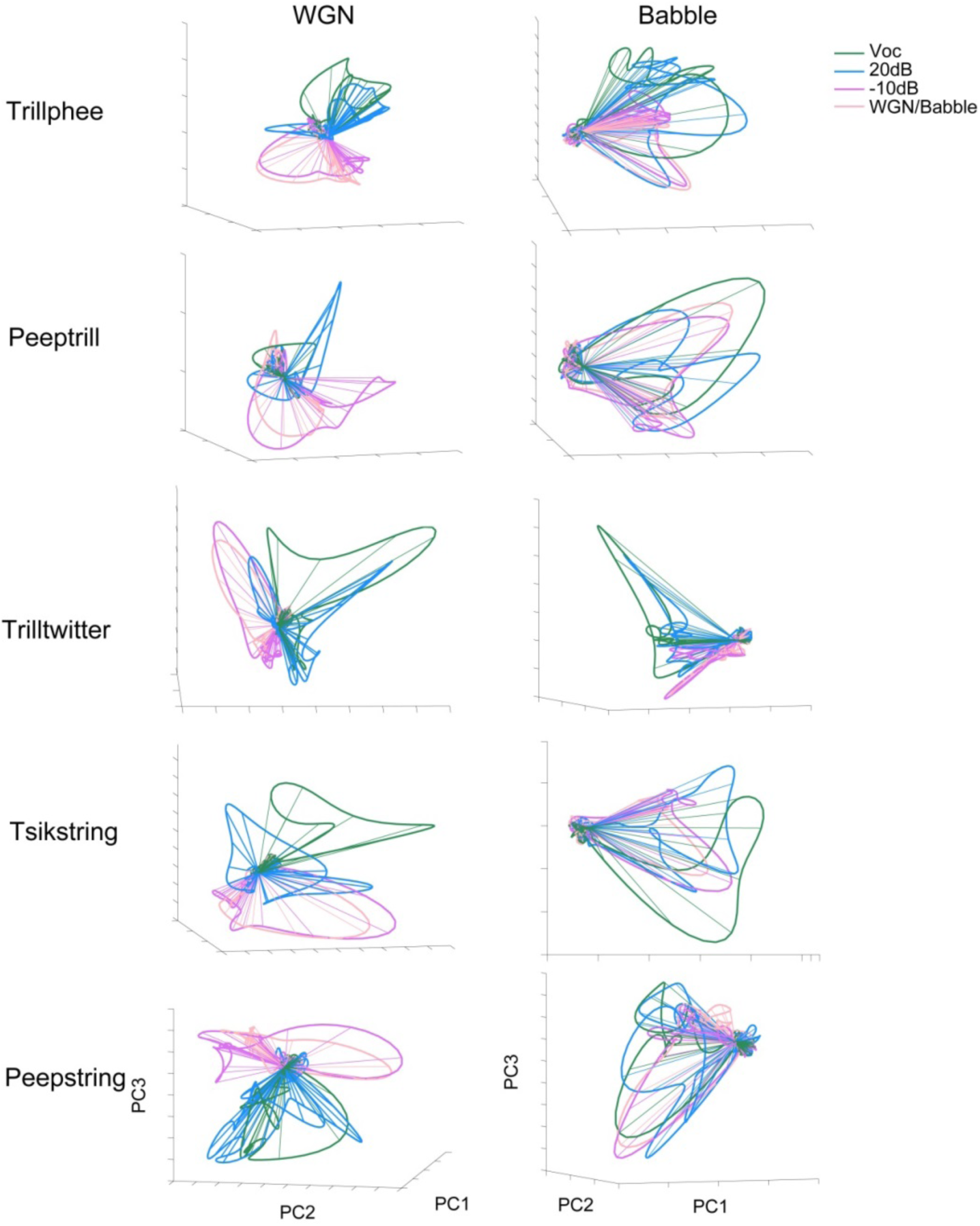
Population-averaged responses to five vocalizations in noise at multiple intensities in 3D response space

### Inter-trajectory angle evolution encodes stimulus SNR

Rotation angles of each trajectory are quantified in Figure 6. Trajectories start with a pre-stimulus portion fluctuating below 60 degrees, greatly increase rotation angles to over 150 degrees following the stimulus onset, and further evolve during stimulus presentation, with particular structures associated with each vocalization. Trajectories at 20 dB SNR share a majority of features with those of clean vocalizations, and trajectories at −10 dB SNR are more noise-like. Again, angle evolution within trajectories of pure vocalizations and pure noise are more separated from each other in the WGN condition than in the Babble condition.

**Figure 6:**
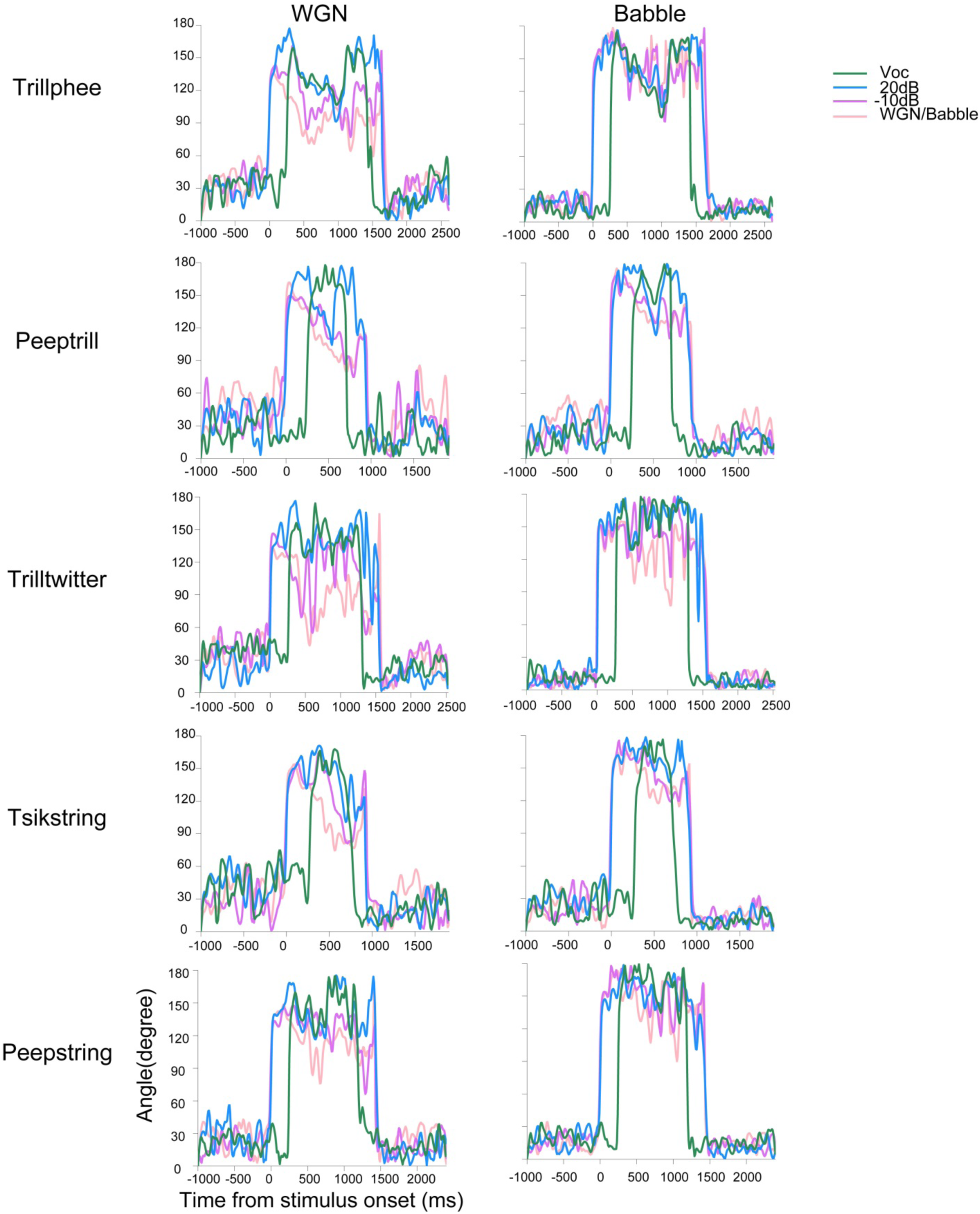
Intra-trajectory angle evolutions in 3D response space for noisy vocalization

To quantify the distance between trajectories, we computed the rotation angles of trajectories of vocalizations at multiple SNRs levels relative to the trajectories of pure vocalizations, with the results shown in Figure 7. Two big peaks indicate the onset and offset responses induced by the two 250-ms noise segments. Time courses between these two peaks show that trajectories at 20 dB SNR have the smallest angular difference from that of pure vocalizations, below 30 degrees. Trajectories of −10 dB SNR and pure noise are further away. Figure 5.11 also quantitatively shows that WGN leads to more separated response trajectories than Babble.

**Figure 7:**
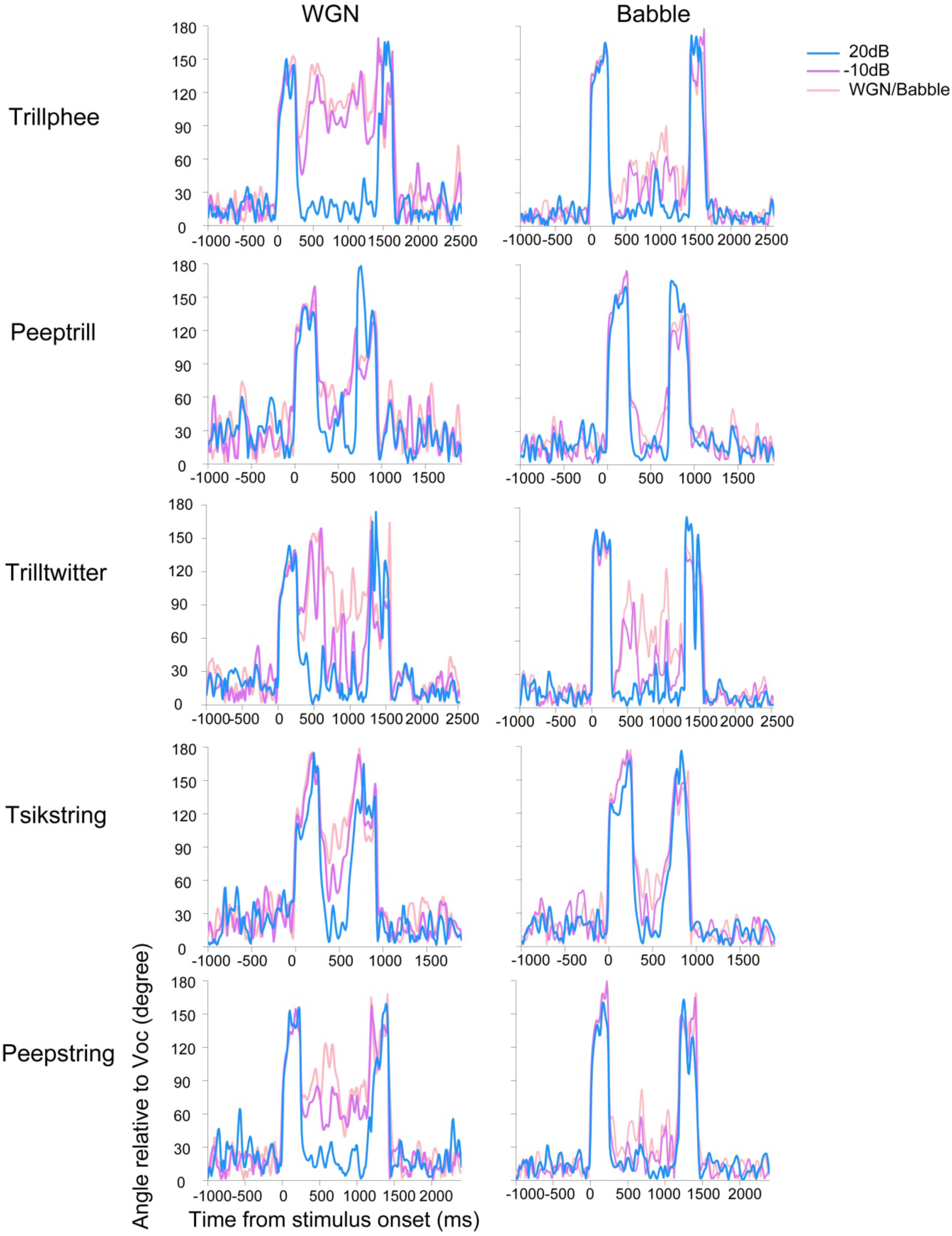
Inter-trajectory angle evolutions in 3D response space for noisy vocalization

### Influence of temporal resolution and temporal integration window on population neural discrimination

In previous analyses, we studied the variability and spatiotemporal structures of population responses to vocalizations at multiple intensities. Population responses averaged over five to ten trials exhibited rich temporal dynamics in terms of rotation angles. Marmoset vocalizations have features spanning a wide range of time scales (Agamaite et al. 2015). Here, we further ask how the trial-by-trial population response discrimination between vocalizations at multiple intensities depends on the temporal resolution. How does the discrimination evolve over time? And how does the number of neurons in the population affect the discrimination?

We quantified the discriminability of population spike trains by building predictive models to decode vocalization types based upon a series of different temporal resolutions, which were used to bin spike trains into response vectors (spike trains of all vocalizations were truncated to the length of the shortest vocalization). As shown in Figure 8, the discrimination accuracy reaches an optimal level at ~ 10 ms, and degrades substantially with widened temporal resolutions. The minimum temporal resolution we tested here is 5 ms, and performance at that level also shows a decreasing trend.

**Figure 8:**
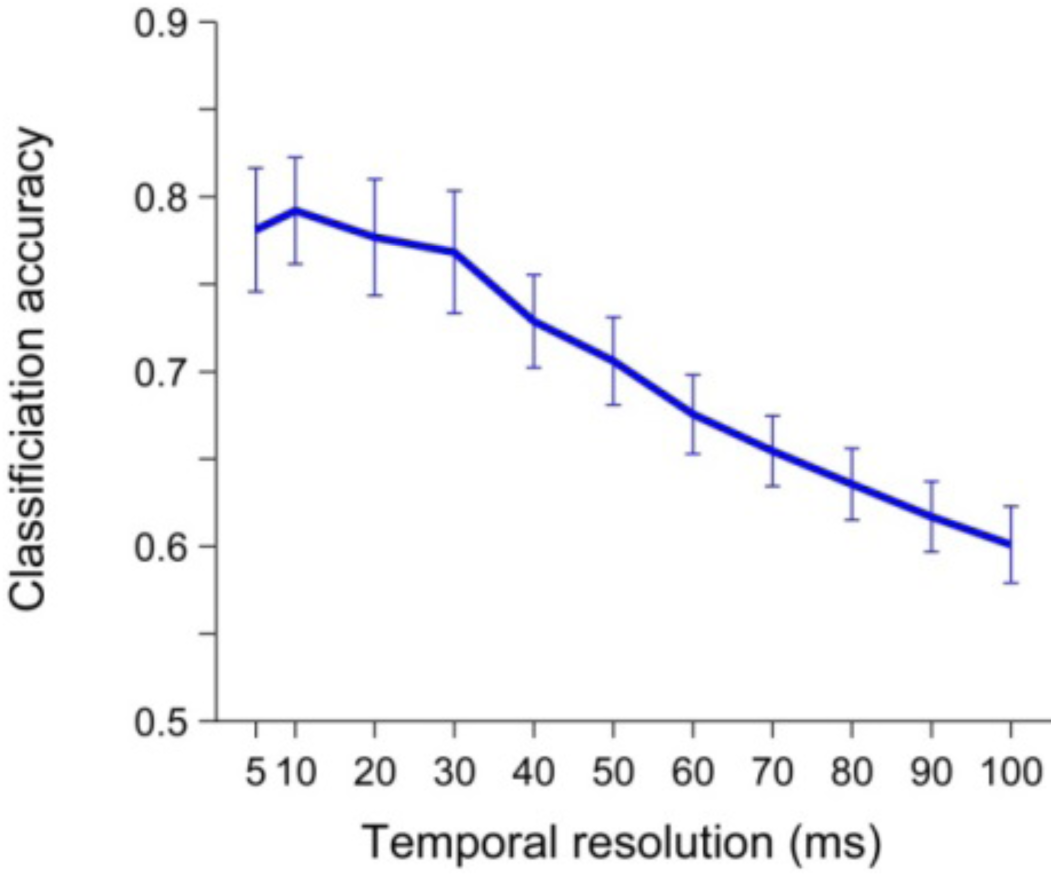
Discriminability of population spike trains in response to vocalizations at a variety of temporal resolutions

To describe the discrimination dynamics over time, we built predictive models with both a single time bin and with increasing numbers of time bins, and obtained the results in Figure 9. Performance of predictive models based upon spontaneous activities is displayed in Figure 9A, along with that of models based upon stimulus–driven activities, as a control. Discrimination with a single time bin begins at the chance level, gradually increases following the onset of vocalizations, and achieves a steady state within 200 ms after stimulus onset. Discrimination with increasing lengths of spike trains is shown in Figure 9B, demonstrating a similar but slightly different trend. It also begins at the chance level, and steadily increases at a relatively fast speed within the first 100 ms after the onset of vocalizations. Later, it enters an oscillating and slowly increasing mode for about 300 ms, and finally reaches a plateau not long before the whole spike train is concluded.

**Figure 9:**
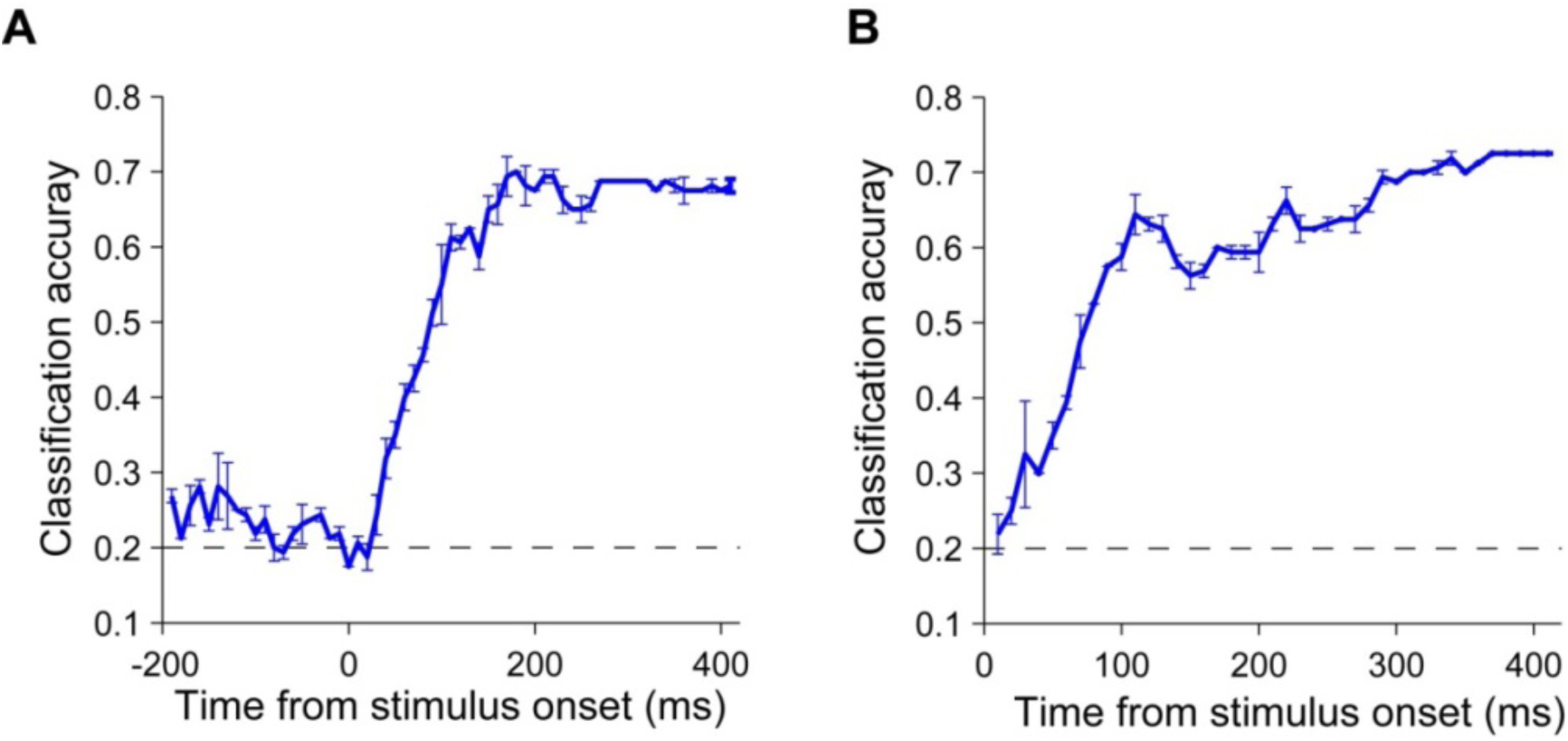
Dynamic discriminability of population spike trains in response to vocalizations over time. (A) Population discrimination of clean vocalizations based on single time bins evolving over time. (B) Population discrimination of clean vocalizations based on increasing length of time bins evolving over time.

Lastly, to evaluate the influence of the number of neurons on discriminability, we randomly sampled various numbers of neurons to build predictive models classifying 20 stimulus labels, until all neurons were included. The resulting discrimination analysis for each vocalization intensity condition is displayed in Figure 10. Generally speaking, as more and more neurons are included as shown in Figure 10A, discrimination improves from the chance level to a plateau when the neuron numbers are between 200 and 300. Neural responses to all vocalizations at 75 dB SPL can be 100% classified when enough neurons are included. The responses at other intensities, however, are not all well classified. In addition, a relative higher intensity does not necessarily guarantee a better performance, as seen by comparing the 55 dB SPL performance of Trillphee and Peeptrill comparing with their 35 dB SPL performance. A better evaluation of the classification performance can be obtained by the confusion matrix in Figure 10B. The confusion matrix provides a clearer view of how likely each neural response to a particular stimulus is to be mistakenly classified as another label. It shows that neural responses to the same vocalization but at different intensities above 15 dB SPL are less likely to be classified as other vocalization types. The matrix demonstrates that the vocalization type is relatively more robustly encoded than the intensity. At 15 dB SPL, misclassified labels seem to be equally distributed among different vocalizations, which makes sense given that the vocalizations are hardly audible at that level.

**Figure 10:**
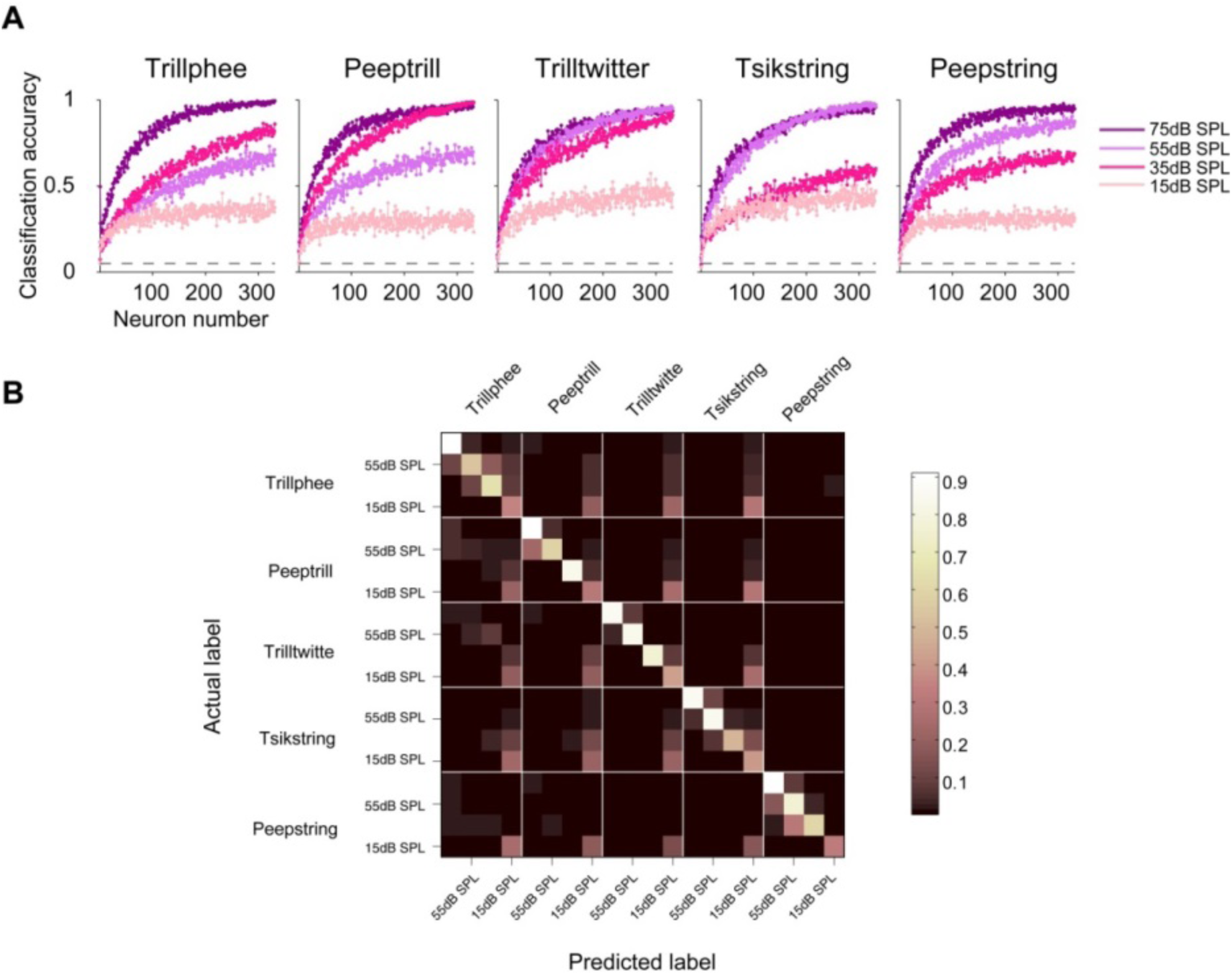
The influence of the number of neurons on population discriminability. (A) Classification accuracy of population responses of five clean vocalizations at different intensities. (B) Confusion matrix of population responses classification.

To evaluate the dependence of discriminability of population responses for vocalizations across multiple SNRs on the temporal resolution, we built predictive models based upon spiking trains binned by time windows of different lengths. Models were built to classify single trial population response to one of the five vocalizations or pure noise (c = 6), and evaluated by the percentage of correctly classified labels. Here, the number of time bins possessed by the shortest vocalization was used. The temporal resolution, in Figure 11, appears to negatively associated with the classification accuracy. Based upon the range of time bins we investigated (5ms ~ 100 ms), a finer temporal resolution appears to provide a better discriminability. In addition, discrimination performance under WGN is about twice that under Babble.

**Figure 11:**
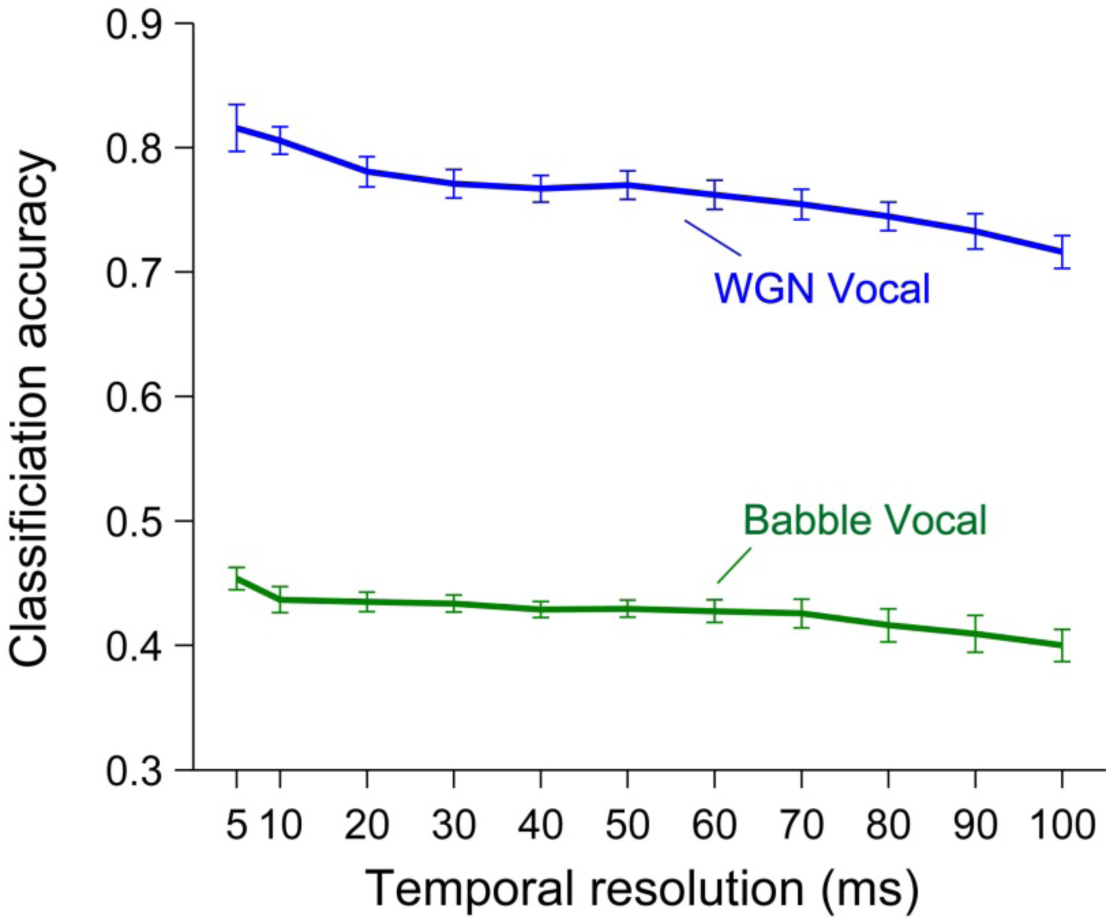
Discriminability of population spike trains between pure noise (WGN/Babble) and noisy vocalizations at a variety of temporal resolutions

More details of the classification performance can be obtained by segregating the accuracy for each vocalization and SNR level as in Figure 12. The performance of each vocalization is displayed as a function of SNR in Figure 12A. For the WGN condition, neural responses to vocalizations delivered with SNR above 5 dB can largely be identified as driven by the correct vocalization type, and neural responses delivered with SNR under −10 dB SNR are most likely to be classified as purely noise-induced, with −5dB and 0dB as the transition points. Neural responses to vocalizations under Babble noise tend to have higher detection thresholds between 0 dB SNR and 10 dB SNR. Under both noise conditions, Peepstring vocalization had the best discrimination over lower SNR levels than other four vocalizations. Whether those wrongly classified neural responses were classified as other types of vocalization of pure noise can be further inferred from Figure 12B. The confusion matrices clearly show that neuron responses driven by a particular vocalization are rarely wrongly classified as other types of vocalizations, except for Tsikstring at −5dB under WGN condition. Noises, instead, exert more interference on the neural responses.

**Figure 12:**
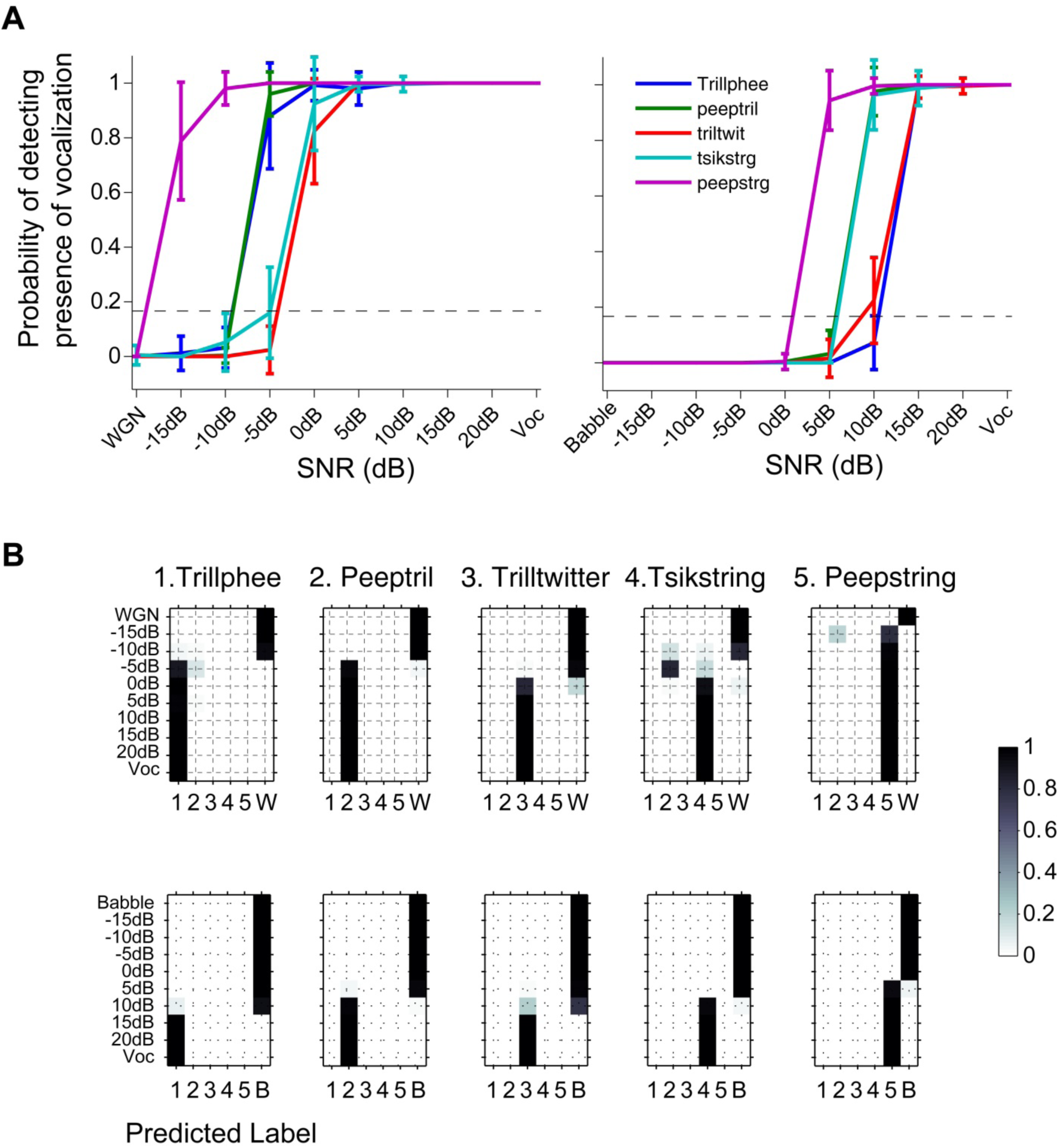
Discriminability of population spike trains between pure noise (WGN/Babble) and noisy vocalizations at a variety of temporal resolutions. (A) Discriminability of population spike trains between pure noise (WGN/Babble) and noisy vocalizations as a function of SNR. (B) Confusion matrices of population response discriminability between pure noise (WGN/Babble) and noisy vocalizations.

We further studied the evolution of population discrimination over time by building predictive models using a single time bin and increased numbers of time bins, as shown in Figure 13. Discrimination based upon a single bin begins at the chance level in Figure 13A, stabilizes at 0.1 for 250 ms of noise preceding the vocalization, and steadily increases following the onset of vocalization in the auditory scene (Figure 1). Babble has a rather low performance based upon single bin response, even below the chance level as shown in Figure 13B. When information was integrated over more and more time bins, the discrimination of population neural responses improved with a steep slope for the first 100 ms following the vocalization onset, and were further boosted under the WGN condition, but reached a plateau under the Babble condition.

**Figure 13:**
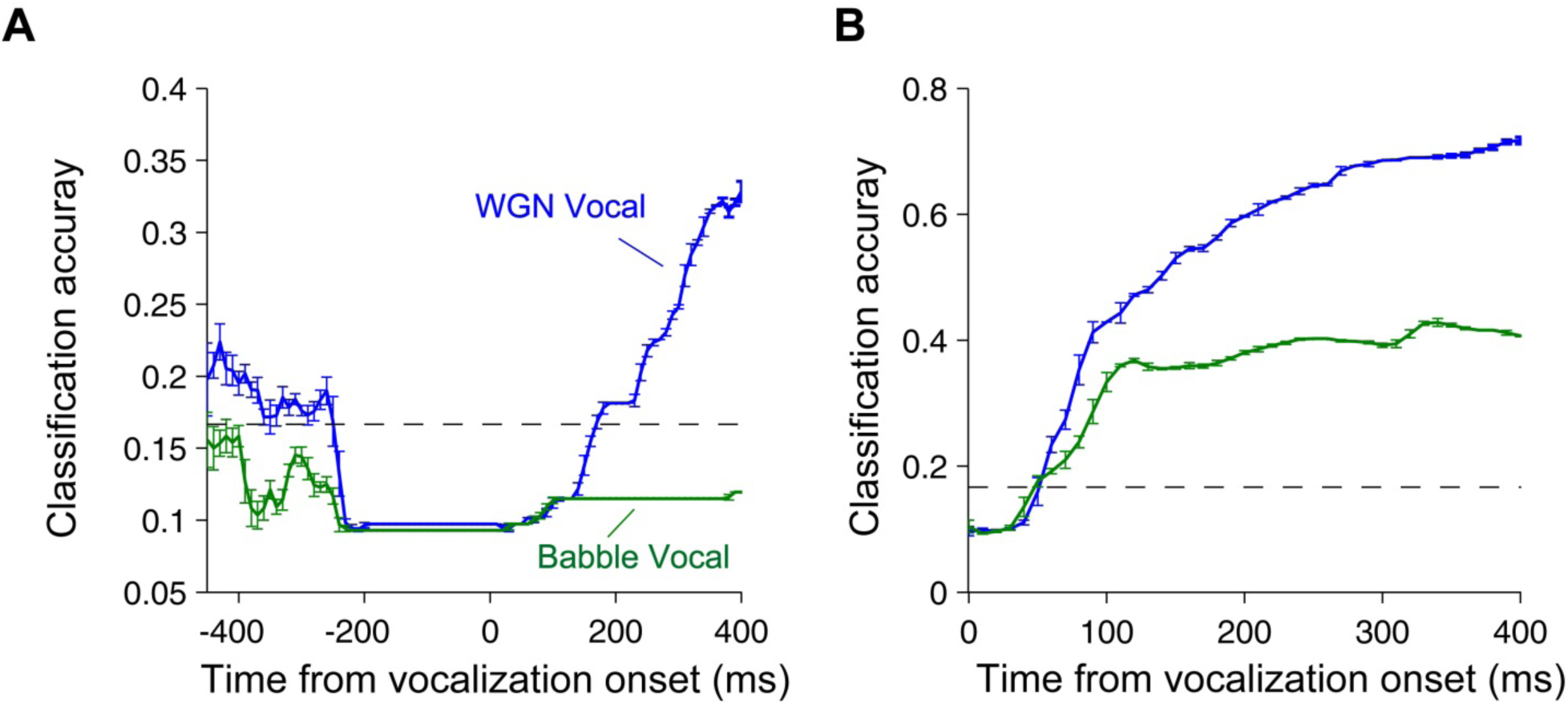
Dynamic discriminability of population spike trains in response to noisy vocalizations over time. (A) Population discrimination of noisy vocalizations based on single time bins evolving over time. (B) Population discrimination of noisy vocalizations based on increasing length of time bins evolving over time.

### Optimal sub-population of neurons with the best discriminative ability

In a previous report, we showed that responses of individual neurons to noisy vocalizations can be categorized into four different groups: *robust*, *balanced*, *insensitive*, and *brittle* (Ni et al. 2016). Here, we investigate the discrimination of subpopulations of neurons by using pure vocalization, 20 dB SNR and pure noise collectively to train the classifiers. Two subpopulations of neurons are shown in Figure 14: the robust group of neurons and population of neurons excluding the brittle group. In Figure 14A, robust groups of neurons generally produce classifiers with a slightly lower detection threshold, but their performance curves are less smoothed. Considering all neurons except the brittle group in Figure 14B adds smoothness and consistency between vocalizations.

**Figure 14:**
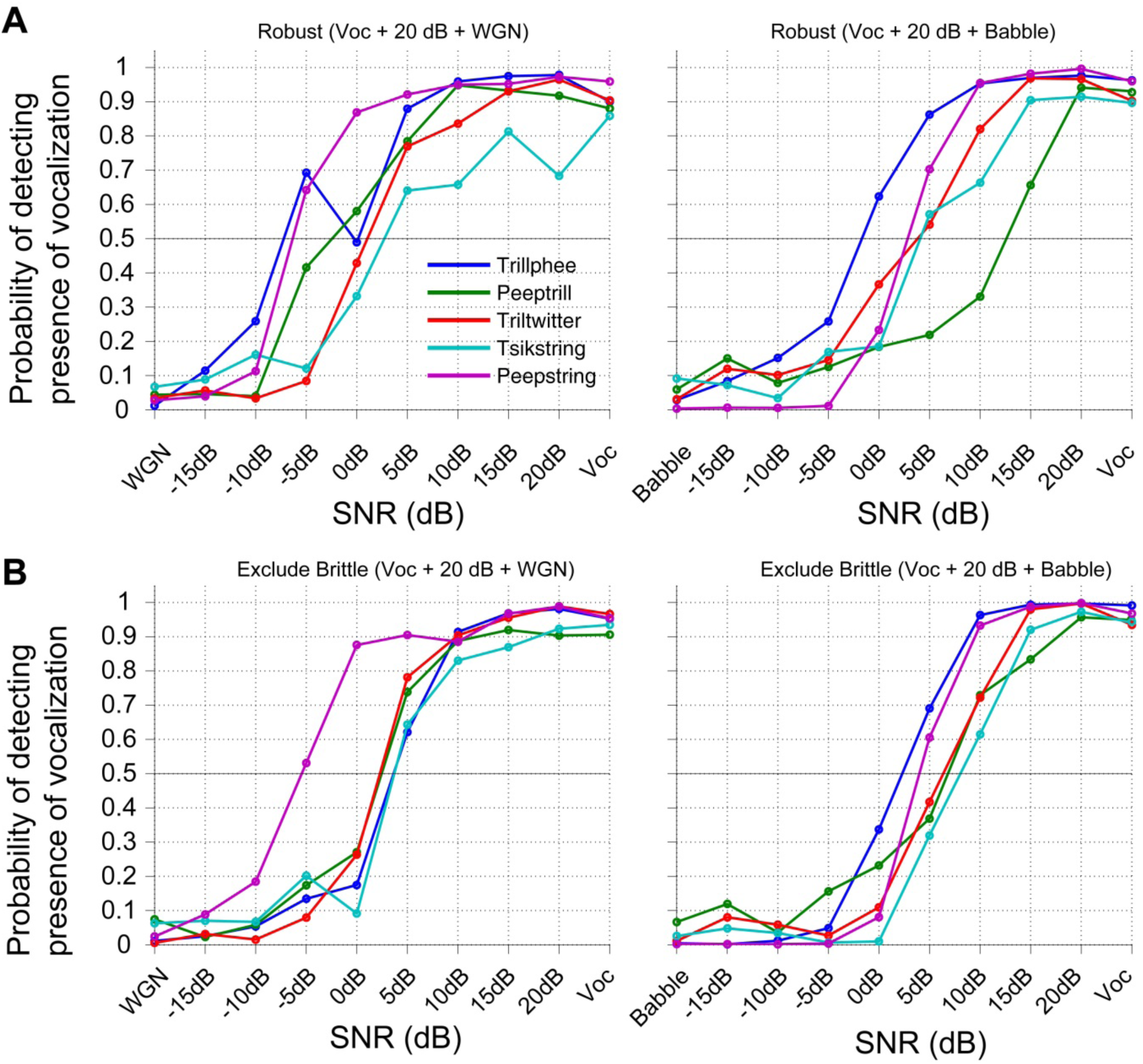
Optimal sub-population for discrimination. (A) Performance of subset of robust neurons discriminating noisy vocalizations. (B)Performance of subset of neurons including robust, balanced, and insensitive to discriminating noisy vocalizations.

### Noise enhances detection threshold

Performances of classifiers heavily depend on the quality of the training dataset. In the machine learning field, it is well known that adding an extra small amount of noise to the training dataset can improve the classifiers’ generalization and obtain better performance (Bishop 1995). Here, we explored the generalization of neural response classifiers by using different training datasets.

For each vocalization, we built separate binary SVM classifiers by using different numbers of time bins, ranging from a single time bin to all the time bins available to that vocalization. Given a trial of population response, the task of the classifiers was to predict whether the response was induced by pure noise or not. The performances of classifiers were averaged over different numbers of time bins. Two groups of training datasets were studied. The first group includes only neural responses to pure noise (labeled as noise) and pure vocalization (labeled as vocalization). The second group includes neural responses to 20 dB SNR as extra training samples labeled as vocalization. The resulting classifier performances are displayed in Figure 15. When only responses to pure noise and pure vocalization are used as training samples, the performances under both noise conditions are not ideal, and greatly degrade around 15 dB SNR as shown in Figure 15A. However, the performance of classifiers trained by the second group of neural responses shows an overall improvement. In Figure 15B, All the lines shift towards the left, with smaller differences between vocalizations, and lead to a lower detection threshold of around 5 dB SNR regardless of noise type. Therefore, by training on neural responses contaminated by a small amount of noise, we can obtain classifiers with more generalized performance over multiple SNR levels.

**Figure 15:**
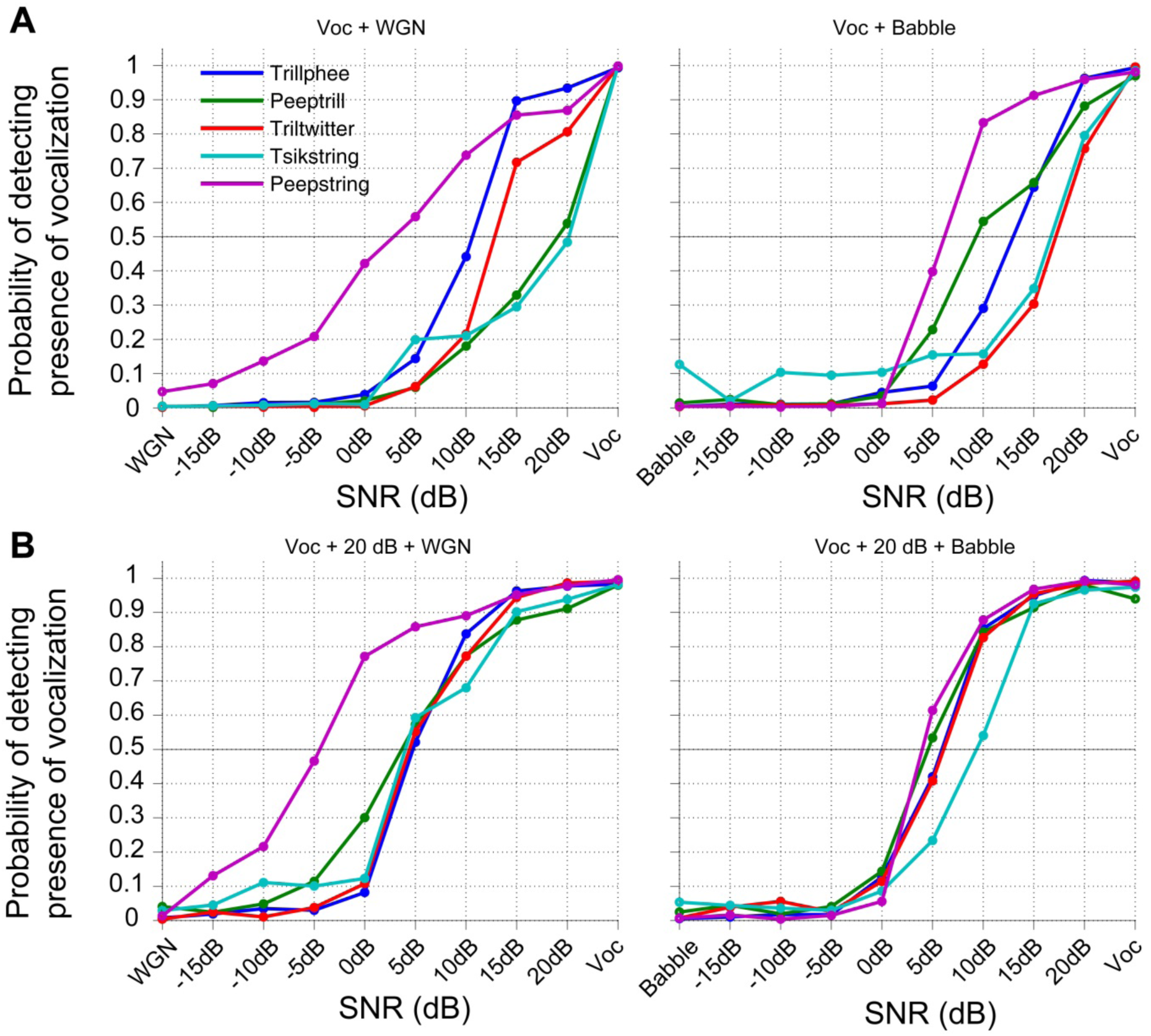
Noise enhances classifiers’ detection threshold. (A) Discriminability of population responses based on vocalizations and pure noises (WGN/Babble). (B) Discriminability of population responses based on vocalizations, 20 dB SNR noisy vocalizations, and pure noises (WGN/Babble).

## Discussion

We examined how the responses of a population of A1 neurons encode vocalizations at multiple intensities and SNR levels by studying the spatiotemporal structures of reduced population responses. We also investigated how well the combined responses of populations of neurons could be used to discriminate among vocalizations under different conditions.

Temporally unstructured stimuli presentations have been demonstrated to induce neural codes of dynamic evolution (Bartho et al. 2009; Friedrich and Laurent 2001; Hegdé and Van Essen 2004; Stopfer et al. 2003; Sugase et al. 1999). By projecting high-dimensional neural responses into a lower dimensional space, we visualized the spatiotemporal structures of population responses induced by complex stimuli: vocalizations at multiple intensities and multiple SNR levels. Even though vocalizations delivered at different intensities are perceptually similar, population responses were progressively differentiated with time, and produced finer discrimination. Different vocalizations can be easily identified by their unique trajectories in space. Some trajectories are relatively smooth and simple, while others are more convoluted, and this variety is associated with the acoustic features of the vocalizations. Differentiation of population response trajectories over time in an auditory scene was dependent on the noise type. WGN noise led to more separable trajectories across SNR levels than Babble, and demonstrated spatiotemporal analysis as a useful indicator of the difficulty of vocalization perception. Consistent with the population coding of tone stimuli (Bartho et al. 2009), population response vectors had the largest rotation during the initial hundreds of milliseconds in response to vocalizations under different conditions, which is probably a common feature shared by population responses to acoustic stimuli regardless of the complexity of stimuli. In addition, we also implemented the same angle evolution analysis using raw population PSTH (data not shown), and revealed a much weaker relationship between the angle evolution and vocalization temporal envelope. This finding indicates that the information of vocalizations is well encoded in a subset of neurons, as the reduced population responses are actually representations of partial covariance in the whole population.

Building neural response classifiers allowed us to investigate the optimal temporal resolution and temporal dynamics of cortical detection and discrimination. Cortical discrimination has been extensively studied for single units, and the optimal temporal resolution was demonstrated to be 10 ms (Machens et al. 2003; Narayan et al. 2006; Rieke 1999; Schneider and Woolley 2010). For our population of neurons, we also found that the temporal resolution for cortical discrimination between vocalizations at multiple intensities on the population level was optimized around 10 ms. This time scale is small enough to capture temporal structures of vocalization, and wide enough to allow integration of information over time, thereby reducing noise. The temporal resolution for cortical discrimination between noisy vocalizations, however, was smaller than 10 ms. In our analysis, the best value was at 5 ms, which is the finest temporal resolution studied here. It is possible that the optimal temporal resolution for noisy vocalization discrimination is below 5 ms. A finer temporal resolution might reduce the interference of the noise component in the auditory scene on vocalization recognition, because a longer time window potentially introduces more noise information, thus confounding the vocalization discrimination. The analysis of the temporal dynamics of discrimination revealed a range for the time scale of integration on the order of hundreds of milliseconds, with ~100 ms for vocalizations at multiple intensities and ~300 ms for noisy vocalizations. The time scale of integration provides information about the speed of accumulation of discrimination accuracy at the population level, and is in similar range to that of single units.

Whether information about sensory stimuli is best represented by the whole population of neurons or a subpopulation of neurons was debated. The discrimination by subpopulations of neurons was particularly studied for noisy vocalizations. We found that the subpopulation of the robust group of neurons and the subpopulations of neurons excluding the brittle group both yielded slightly lower detection thresholds than the whole population, thus a better discrimination performance. This result indicates that the brittle group of neurons contributes as a neural distractor for noisy vocalization discrimination, and that information about vocalization is better encoded by a subpopulation of neurons instead. We also built classifiers to demonstrate that we can generalize the discrimination over lower SNR levels by using neural responses contaminated by a little acoustic noise as training samples. While we are not saying that the brain actually decodes vocalization information in the same way that our classifiers do, the results are consistent with a previous psychoacoustic study that demonstrated that introducing weak noises in perception improved the detection thresholds of target signals (Zeng et al. 2000).

In summary, we investigated our data with population analytic techniques and revealed population response dynamics that cannot be fully evaluated by single-unit analysis alone.

The authors declare no competing financial interests.

## Acknowledgment

This work was supported by the National Institutes of Health grant R01-DC009215. We thank Kim Kocher for valuable assistance with animal training and data collection. Our thanks also go to Wensheng Sun for her help with neurophysiology experiment preparation.

